# Landscape-level DNA metabarcoding study in the Pannonian forests reveals differential effects of slope aspect on taxonomic and functional groups of fungi

**DOI:** 10.1101/281337

**Authors:** József Geml

## Abstract

In temperate regions, slope aspect is one of the most influential drivers of environmental conditions at landscape level. The effect of aspect on vegetation has been well studied, but virtually nothing is known about how fungal communities are shaped by aspect-driven environmental conditions. I carried out DNA metabarcoding of fungi from soil samples taken in a selected study area of Pannonian forests to compare richness and community composition of taxonomic and functional groups of fungi between slopes of predominantly southerly vs. northerly aspect and to assess the influence of selected environmental variables on fungal community composition. The deep sequence data presented here (i.e. 980 766 quality-filtered sequences) indicate that both niche (environmental filtering) and neutral (stochastic) processes shape fungal community composition at landscape level. Fungal community composition correlated strongly with aspect, with many fungi showing preference for either south-facing or north-facing slopes. Several taxonomic and functional groups showed significant differences in richness between north-and south-facing slopes and strong compositional differences were observed in all functional groups. The effect of aspect on fungal communities likely is mediated through contrasting mesoclimatic conditions, that in turn influence edaphic processes as well as vegetation. Finally, the data presented here provide an unprecedented insight into the diversity and landscape-level community dynamics of fungi in the Pannonian forests.

## INTRODUCTION

In temperate regions, slope aspect is among the most influential factors that drive the physical environment at the landscape level. The aspect effect is driven primarily by the variation in the amount of solar radiation per surface area, which is a function of the angle of incidence of solar radiation. As a result, slopes oriented toward the Equator receive higher intensity and greater duration of solar radiation, which can be 50% higher on south-facing than on north-facing slopes in the northern hemisphere (Geiger 1965; Rosenberg *et al*. 1983; Gilliam *et al*. 2014). These contrasting energy inputs can profoundly alter mesoclimatic conditions, particularly air and upper soil temperature, which, in turn, affect relative humidity, evapotranspiration, soil moisture and edaphic processes (Fekedulegn *et al*. 2003). As a result, slopes with poleward (in this case, northerly) aspect generally are cool and more humid, while slopes with southerly aspect tend to be markedly warmer and drier, particularly in mid-and high latitudes (Holland & Steyn 1975; Méndez-Toribio *et al*. 2016).

Such aspect-related contrasts in the abiotic environment are intuitively expected to have influence on the composition of biotic communities. Indeed, the effects of aspect on vegetation are well known and clear compositional differences have been found between north-and south-facing slopes in various ecosystems, e.g., in temperate and Mediterranean forests (Whittaker 1956; Sternberg & Shoshany 2001; Gilliam *et al*. 2014), temperate grasslands (Schmidt 2013), boreal forests (Hollingsworth *et al*. 2006), and arctic tundra (Walker *et al*. 1994). On the other hand, virtually nothing is known about the influence of aspect on richness and community composition of fungi at landscape level, except two recent studies focusing on arbuscular mycorrhizal fungal richness in boreal forests and arid steppes in China (Chu *et al*. 2016; Liu *et al*. 2017). Therefore, how slope aspect affects richness and community composition of various taxonomic and functional groups of fungi remains unknown.

According to the macroecological study of Tedersoo *et al*. (2014), global-scale fungal diversity and distribution patterns are primarily influenced by climatic factors, mainly mean annual temperature and precipitation, followed by edaphic factors, particularly pH, and spatial patterns due to dispersal limitation. On the other hand, the coupling between vegetation and soil fungal community composition and richness appears to be much weaker, with the exception of ectomycorrhizal (ECM) fungi (Tedersoo *et al*. 2014; Peay *et al.* 2016). Therefore, it is reasonable to hypothesize that the above-mentioned aspect-driven environmental differences are expected to influence the diversity and distribution of fungal communities at landscape level as well.

In this study, I tested the effects of aspect on the richness and composition of taxonomic and functional groups of fungi at a selected study site in the Pannonian forests, where regional climatic conditions and geological parent material were uniform in order to minimize confounding factors. The Pannonian biogeographical region represents unique ecosystems in Europe (Sundseth 2009). It covers the inner parts of the Carpathian basin and is characterized by a great wealth of floristic and faunistic elements from different parts of Eurasia. More specifically, in addition to the broadly distributed Eurasian species, there are numerous sub-Mediterranean, continental, Pontic and Balkanian species. The distinctiveness of the Pannonian biogeographic region comes from the combination of these elements, as well as the occurrence of Pannonian endemics (Fekete *et al.* 2016). The geologically diverse mountain range of the Északi-középhegység (North Hungarian Mountains) represents the northern part of the Pannonian biogeographic region. The region is characterized by high habitat diversity, partly due to the complex geology which features a wide variety of calcareous, volcanic and igneous rocks (Pelikán 2010), and partly due to the diverse topography that creates a broad spectrum of mesoclimatic conditions. This diversity of habitats allows for the coexistence of sub-Mediterranean, continental, Atlantic, and Carpathian floristic and faunistic elements often in close proximity, depending on slope, aspect, elevation, and geological parent material (Suba 1983; Vojtkó 2002; Vojtkó *et al*. 2010). With respect to macrofungi, there is a long history of sporocarp-based studies in various areas of the Északi-középhegység (Bohus & Babos 1960; Takács & Siller 1980; Rimóczi 1992, 1994; Tóth 1999; Siller *et al*. 2002; Albert & Dima 2005; Egri 2007; Pál-Fám *et al*. 2007; Rudolf *et al*. 2008; Siller 2010; Siller & Dima 2014). In addition, morphological and molecular analyses of roots colonized by ECM genera *Humaria*, *Gevea*, *Tomentella*, and *Tuber* have been carried out in a well-preserved montane beech forest reserve (Kovács & Jakucs 2006; Erős-Honti *et al*. 2008; Jakucs *et al*. 2015). However, the diversity and distribution of fungi, particularly microscopic fungi, in various habitats types are still unexplored and I am not aware of any molecular study assessing the local (alpha) diversity and/or community turnover (beta diversity) of fungal communities in the Pannonian forests.

The objectives of this DNA metabarcoding study were 1) to provide the first insights into the kingdom-wide taxonomic and functional diversity of fungi in different Pannonian forest types on a landscape scale; 2) to compare richness and community composition of fungi among sampling sites among predominantly south-and north-facing slopes; and 3) to assess the influence of selected environmental variables on fungal community composition.

## MATERIALS AND METHODS

### Study area

Within the broader region of the Északi-középhegység characterized above, the focal area of this study is located in the valley of the Tarna creek situated between the Mátra and the Bükk, the two highest mountains in Hungary. The study area is characterized by rolling hills of low elevations (200-500 m a.s.l.) that are mostly covered with managed secondary forests. The regional climate is subcontinental, with warm summers (mean July temperature ca. 20°C), cold winters (mean January temperature ca. -3°C), and mean annual temperature and precipitation of ca. 9.5°C and ca. 580 mm, respectively (Tóth 1983; Horváth & Gaálová 2007).

Pannonian-Balkanic turkey oak-sessile oak forests (*Quercetum petraeae-cerris*, Natura 2000 code 91M0) form the zonal vegetation type on gentle slopes and in placor settings in the study area as well as in other colline (200-450 m a.s.l.) areas of the Északi-középhegység. They are dominated by oaks, such as *Quercus dalechampii*, *Q. petraea*, and *Q. cerris*, and can attain 15-25 m canopy height. Within the turkey oak-sessile oak zone, Pannonian thermophilous downy oak forests (*Corno-Quercetum pubescentis*, 91H0) generally develop on shallow and rocky soil on steep (> 20°), south-facing slopes. These xerothermic forests are also dominated by the same oak species as the zonal forest type, but the canopy remains low or medium tall (8-10 m), with the characteristic appearance of *Q. pubescens*, particularly in the western half of Hungary, where this vegetation type is more widespread. In the Északi-középhegység, thermophilous oak forests tend to occur in isolated patches with particularly warm and dry mesoclimate (Vojtkó 2002; Borhidi 2003; Vojtkó *et al*. 2010; Bölöni *et al.* 2011).

North-facing, gentle slopes are mostly covered by Pannonian sessile oak-hornbeam forests (*Carici pilosae-Carpinetum*, 91G0), that are predominantly mesophilous, submontane (400-600 m a.s.l.) forests, characteristic of cool, relatively humid habitats on deep soils, with a canopy height of 25-30 m. These mixed forests are dominated by *Carpinus betulus* and *Quercus petraea*, with *Acer campestre*, *A. platanoides*, *Cerasus avium*, and *Tilia cordata* occurring sporadically. In the low-elevation landscape of the study area, extrazonal submontane beech forests (*Melittio-Fagetum*) occur very sporadically and are restricted to steep, north-facing slopes with the coldest mesoclimatic conditions. These are tall (25-35 m), mesic forests, where *Fagus sylvatica* generally is monodominant. In zonal settings in the Északi-középhegység, beech forests (9130) are represented by submontane beech forest (*Melittio-Fagetum*) between 400 and 750 m a.s.l. and montane beech forest (*Aconito-Fagetum*) above 750 m a.s.l., the latter featuring many species with Carpathian distributions (Borhidi 2003; Vojtkó *et al.* 2010; Bölöni *et al*. 2011).

The fourteen sampling sites were evenly distributed in two hills (Kis-várhegy and Nagy-várhegy) just south of the village of Sirok, Heves county, Hungary (Fig. 1, Table 1). The location was chosen because it features, in an unusually small area (< 2 km^2^), the above-mentioned major Pannonian forest types of northern Hungary, which are distributed according to aspect-based differences in mesoclimatic conditions in a not spatially clustered manner. Furthermore, potentially confounding factors that often limit the interpretation of ecological studies in natural habitats are minimal: 1) the Kis-várhegy and Nagy-várhegy are uniformly composed of Jurassic calcareous limestone (Pelikán 2010); 2) the regional climate is uniform throughout the small study area; 3) all forest types are represented by old-growth (> 80y) secondary forests; 4) the elevation differences among the sampling sites are minor (< 60 m) and are not related to aspect or to any other environmental conditions; and 5) the spatial proximity and the continuous natural or semi-natural landscape are expected to facilitate unrestricted dispersal of fungal propagules among the sites by wind, water or by wildlife. Therefore, I hypothesized that any observed compositional differences among the samples would be the result of niche-based and stochastic processes, with the former primarily referring to environmental filtering according to differences in abiotic and biotic factors driven by topography: predominantly northerly vs. southerly aspect and to a lesser extent by slope angle.

**Table 1.** Sampling sites included in this study with code, locality, forest type, dominant tree species, slope aspect, northerly aspect expressed in degrees (south=0, north=180), slope angle, and geographic coordinates. Abbreviations for tree genera are: *C.*: *Carpinus* (Betulaceae), *F.*: *Fagus* (Fagaceae), and *Q*.: *Quercus* (Fagaceae). Locations are displayed in a map in Figure 1.

| Site code | Locality | Forest type | Dominant tree species | Aspect | Northerly aspect<br>(degrees) | Slope angle<br>(degrees) | Latitude<br>(degrees) | Longitude<br>(degrees) |
| --- | --- | --- | --- | --- | --- | --- | --- | --- |
| KV1 | Kis-várhegy | *Quercetum petraeae-cerris* | *Q. dalechampii*, *Q. petraea*, *Q. cerris* | SW | 13 | 22.85 | 47.918885 | 20.187345 |
| KV2 | Kis-várhegy | *Corno-Quercetum pubescentis* | *Q. dalechampii*, *Q. petraea*, *Q. cerris* | S | 5 | 5.71 | 47.918840 | 20.188193 |
| KV3 | Kis-várhegy | *Corno-Quercetum pubescentis* | *Q. dalechampii*, *Q. petraea*, *Q. cerris* | S | 2 | 29.71 | 47.919033 | 20.190292 |
| KV4 | Kis-várhegy | *Corno-Quercetum pubescentis* | *Q. dalechampii*, *Q. petraea*, *Q. cerris* | S | 10 | 24 | 47.918927 | 20.192524 |
| KV5 | Kis-várhegy | *Carici pilosae-Carpinetum* | *C. betulus*, *Q. petraea* | N | 175 | 19.43 | 47.921958 | 20.193339 |
| KV6 | Kis-várhegy | *Carici pilosae-Carpinetum* | *C. betulus*, *Q. petraea* | N | 170 | 22.86 | 47.921977 | 20.192281 |
| KV7 | Kis-várhegy | *Carici pilosae-Carpinetum* | *C. betulus*, *Q. petraea* | N | 165 | 25.14 | 47.921792 | 20.191034 |
| NV1 | Nagy-várhegy | *Carici pilosae-Carpinetum* | *C. betulus*, *Q. petraea* | WNW | 115 | 17.14 | 47.912658 | 20.181935 |
| NV2 | Nagy-várhegy | *Carici pilosae-Carpinetum* | *C. betulus*, *Q. petraea* | NW | 145 | 33.14 | 47.916270 | 20.184321 |
| NV3 | Nagy-várhegy | *Melittio-Fagetum* | *F. sylvatica* | N | 178 | 37.71 | 47.917315 | 20.186580 |
| NV4 | Nagy-várhegy | *Melittio-Fagetum* | *F. sylvatica* | N | 179 | 38.85 | 47.917191 | 20.188443 |
| NV5 | Nagy-várhegy | *Quercetum petraeae-cerris* | *Q. dalechampii*, *Q. petraea*, *Q. cerris* | WNW | 115 | 22.85 | 47.913477 | 20.183917 |
| NV6 | Nagy-várhegy | *Corno-Quercetum pubescentis* | *Q. dalechampii*, *Q. petraea*, *Q. cerris* | SW | 15 | 24.28 | 47.906740 | 20.180339 |
| NV7 | Nagy-várhegy | *Quercetum petraeae-cerris* | *Q. dalechampii*, *Q. petraea*, *Q. cerris* | SW | 18 | 8 | 47.906650 | 20.181740 |

**Fig. 1.**
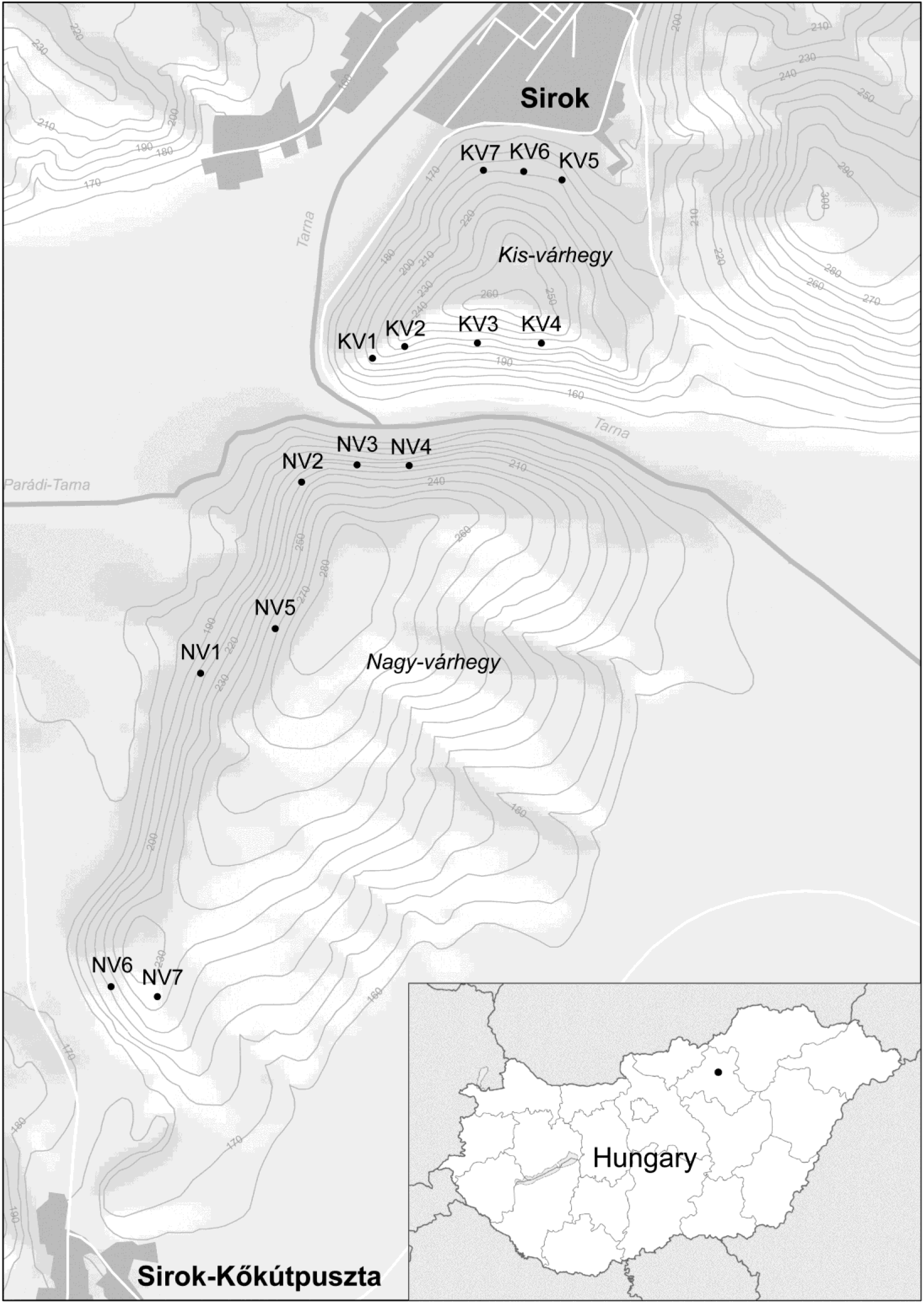
A map of the sampling localities, with the location of the region of study in Hungary (inset). Full names, vegetation types, topographic variables, and geographic coordinates corresponding to the sampling localities are listed in Table 1.

### Sampling and molecular work

For this study, fourteen sites were chosen that represented forests of northerly and southerly aspects on both the Kis-and Nagy-várhegy. At each site (ca. 10 × 25 m), 20 samples (5 cm^3^ each) of top soil from underneath the litter layer were taken at each sampling site in such a way that samples were at least 2 m from each other to minimize the probability of sampling the same genet repeatedly. Soil samples collected at a given site were pooled, resulting in a composite soil sample for each site. Ca. 20 g of each composite sample was kept frozen until DNA extraction, while the rest was used for soil chemical analyses to measure pH (water-based), and total carbon (C) and nitrogen (N) contents following Sparks *et al*. (1996).

Genomic DNA was extracted from 0.5 ml of soil from each composite sample using NucleoSpin^®^ soil kit (Macherey-Nagel Gmbh & Co., Düren, Germany), according to manufacturer’s protocol. The ITS2 region (ca. 250 bp) of the nuclear ribosomal DNA repeat was PCR amplified using primers fITS7 (Ihrmark *et al.,* 2012) and ITS4 (White *et al.,* 1990). The ITS4 primer was labelled with sample-specific Multiplex Identification DNA-tags (MIDs). The amplicon library was sequenced at Naturalis Biodiversity Center (Naturalis) using an Ion 318™ Chip and an Ion Torrent Personal Genome Machine (Life Technologies, Guilford, CT, U.S.A.). Detailed protocols of the molecular work are described in Geml *et al.* (2014b).

### Bioinformatic work

The initial clean-up of the raw data was carried out using Galaxy (https://main.g2.bx.psu.edu/root), in which the sequences were sorted according to samples and adapters (identification tags) were removed. The primers were removed and poor-quality ends were trimmed off based on 0.02 error probability limit in Geneious Pro 8 (BioMatters, New Zealand). Subsequently, sequences were filtered using USEARCH v.8.0 (Edgar, 2010) based on the following settings: all sequences were truncated to 200 bp and sequences with expected error > 1 were discarded. For each sample, sequences were collapsed into unique sequence types, while preserving their counts. The quality-filtered sequences from all samples were grouped into operational taxonomic units (OTUs) at 97% sequence similarity and putative chimeric sequences were removed using USEARCH. I assigned sequences to taxonomic groups based on pairwise similarity searches against the curated UNITE+INSD fungal ITS sequence database (version released on October 10, 2017), containing identified fungal sequences with assignments to Species Hypothesis (SH) groups delimited based on dynamic sequence similarity thresholds (Kõljalg *et al*. 2013). After excluding OTUs with < 80% similarity or < 150 bp pairwise alignment length to a fungal sequence, 6216 fungal OTUs were retained, representing a total of 980 766 quality-filtered sequences, including 1930 global singletons. Global singletons are routinely excluded from metabarcoding studies, because the vast majority of them represent erroneous sequences (Lindahl *et al*. 2013; Geml *et al*. 2016, 2017). However, this practice likely results in the exclusion of some real high-quality data on locally rare species. Because, to my best knowledge, this is the first metabarcoding study in the Pannonian forests, I intended to keep all high-quality data, including 343 singletons with > 98% sequence similarity to a reference fungal SH, a conservative threshold for conspecificity in most fungal groups (Kõljalg *et al.* 2013), and excluded the remaining 1587 singletons from further analyses. DNA sequences have been deposited in NCBI (accession numbers provided upon manuscript acceptance).

Only OTUs with > 90% similarity to a fungal SH with known ecological function were assigned to one of the following functional groups: animal pathogens, ECM fungi, lichens, litter decomposers, mycoparasites, plant pathogens, root-associated fungi (non-ECM orchid and ericoid mycorrhizal fungi and root endophytes), saprotrophs (generalists), and wood decomposers. Arbuscular mycorrhizal fungi were not included in the analyses, because only one representative OTU was found in the quality-filtered dataset. The initial functional assignments were made by FunGuild (Nguyen *et al*. 2015) and were manually checked afterwards. For genera that are known to comprise species from multiple functional guilds (e.g., *Amanita*, *Entoloma*, *Ramaria*, and many SH groups in the Sebacinales), I assigned ecological function for each OTU individually, based on available ecological information for the matching SH in the UNITE database.

### Statistical analyses

I normalized the OTU table for subsequent statistical analyses by rarefying the number of high-quality fungal sequences to the smallest library size (36 946 reads). The resulting matrix contained 4312 fungal OTUs. Linear regression analyses in R (R Development Core Team 2013) were used to examine relationships between aspect and richness of taxonomic and functional groups of fungi as well as between aspect and environmental variables, i.e. edaphic factors (pH, C, N, and C/N) and the relative abundance values of the two dominant tree genera (*Carpinus* and *Quercus*) that were distributed throughout the majority of sites. When aspect is treated as a continuous variable from 0° to 360°, the two extreme values of this interval refer to the same aspect in the landscape (north). Therefore, aspect was expressed as northerly aspect in degrees (south: 0°, north: 180°) following Calef *et al*. (2005), which better reflects the well-known ecological differences between north-and south-facing slopes.

I used the *vegan* R package (Oksanen *et al*. 2015) to run non-metric multidimensional scaling (NMDS) on the Hellinger-transformed OTU table and a secondary matrix containing environmental variables, which were standardized using the *scale* function in R. Ordinations were run separately for functional groups as well as for all fungi with the following specifications: distance measure = Bray-Curtis, dimensions = 2, initial configurations = 100, model = global, maximum number of iterations = 200, convergence ratio for stress = 0.999999. I used the *envfit* R function to fit the above-mentioned environmental variables and richness of various taxonomic and functional groups onto the NMDS ordinations and plotted isolines of northerly aspect on the NMDS ordinations using the *ordisurf* function. In addition, I tested whether fungal communities were statistically different among forest types using the multiresponse permutation procedure (MRPP) and determined any preferences of individual OTUs for forest types and for north-or south-facing slopes using indicator species analyses (Dufrêne & Legendre 1997) in PC-ORD v. 6.0 (McCune & Grace 2002).

A series of mantel tests were carried out in PC-ORD to reveal any spatial autocorrelation in environmental variables as well as fungal community composition among the sampling sites and to measure correlation between fungal community composition and environmental variables. Furthermore, a series of partial mantel tests were applied to differentiate the effects of northerly aspect, tree genera, and edaphic factors on fungal community structure. Finally, I estimated what proportions of the total variation in fungal community composition were explained by topography, tree genera, and edaphic factors using variation partitioning (Borcard *et al*. 1992; Legendre *et al.* 2005) with a series of redundancy analyses (RDAs) in Canoco 5 (Microcomputer Power, Ithaca, NY, USA). The expectation in purely neutral (stochastically assembled) communities is no correlation between the environmental distance and the community dissimilarity of two sites (Smith & Lundholm 2010).

## RESULTS

### Correlation of elevation with fungal richness and environmental variables

In total, the quality-filtered and rarefied dataset contained 4312 fungal OTUs in the 14 soil samples taken from the representative types of Pannonian forests. Of these, 2196 OTUs had > 90% similarity to a fungal SH with known ecological function were assigned to functional groups. Richness values of most functional groups showed significant (*p* < 0.05) or marginally (*p* < 0.1) significant correlation with northerly aspect. For example, I found positive correlation in root-associated fungi (*r*^2^ = 0.231; *p* = 0.047), mycoparasites (*r*^2^ = 0.148; *p* = 0.096), and wood decomposers (*r*^2^ = 0.204; *p* = 0.059) with northerly aspect, while negative correlation was observed in lichens (*r*^2^ = 0.154; *p* = 0.091), plant pathogens (*r*^2^ = 0.231; *p* = 0.047), and generalist saprotrophs (*r*^2^ = 0.615; *p* < 0.001) (Fig. 2). Richness of several taxonomic groups also correlated with northerly aspect with varying significance, positively in Agaricomycetes (*r*^2^ = 0.261; *p* = 0.036), Tremellomycetes (*r*^2^ = 0.211; *p* = 0.056), Leotiomycetes (*r*^2^ = 0.166; *p* = 0.083), and Mortierellomycota and Mucoromycota that had formerly been classified to Zygomycota (*r*^2^ = 0.465; *p* = 0.004), and negatively in Dothideomycetes (*r*^2^ = 0.605; *p* < 0.001), Eurotiomycetes (*r*^2^ = 0.523; *p* = 0.002), and Sordariomycetes (*r*^2^ = 0.386; *p* = 0.011) (Fig. 2). Among the measured edaphic factors, only pH correlated significantly with northerly aspect, showing strong negative correlation (*r*^2^ = 0.516; *p* = 0.002). With respect to vegetation, the relative abundance of *Quercus* correlated negatively (*r*^2^ = 0.837; *p* < 0.001), while that of *Carpinus* correlated positively (*r*^2^ = 0.269; *p* = 0.033) with northerly aspect (Fig. 2).

**Fig. 2.**
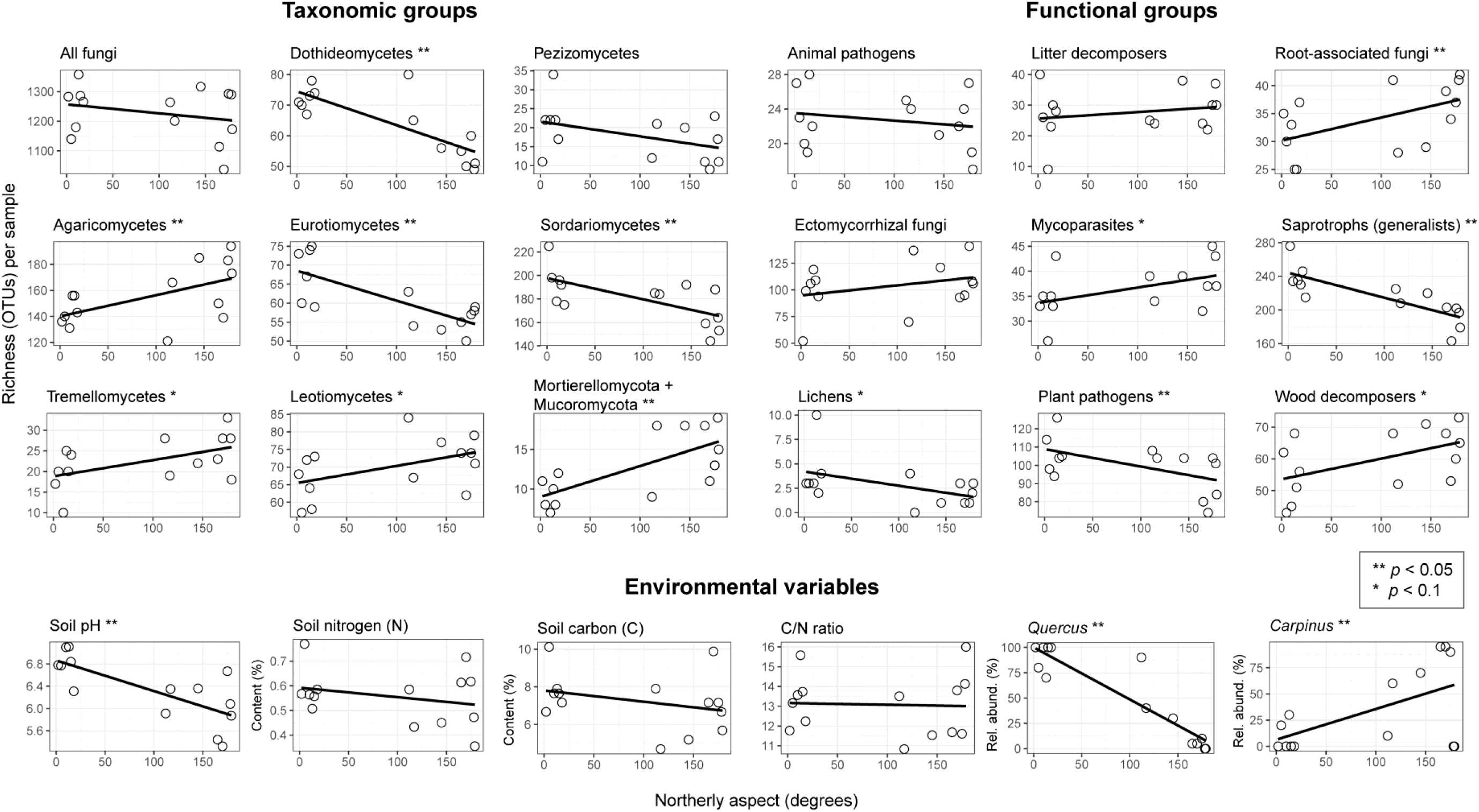
Correlations between northerly aspect and operational taxonomic unit (OTU) richness of functional and taxonomic groups as well as environmental variables explored using linear regression. Significant (*p* < 0.05) and marginally significant (*p* < 0.1) correlations are marked by ** and *, respectively.

### Comparing fungal community composition among the sampling sites

All NMDS analyses resulted in 2-dimensional solutions with the following final stress values for all fungi (0.0693), animal pathogens (0.1396), ECM fungi (0.1046), litter fungi (0.1575), mycoparasites (0.0871), plant pathogens (0.0679), root-associated fungi (0.1342), saprotrophs (0.0947), and wood decomposers (0.1471). In all cases, the NMDS ordinations revealed strong structuring of fungal communities according to aspect. In addition, a weaker, but often clear separation could be observed among the four forest types. Community composition of all fungi in the sampling sites clearly structured according aspect which represented the first axis (*r* = 0.999) (Fig. 3). Soil pH (*r* = -0.993) correlated strongly with the first axis in that south-facing slopes had higher pH than north-facing ones. With respect to trees, relative abundance of oaks (*r* = -0.977) correlated negatively, while both hornbeam (*r* = 0.587) and beech (*r* = 0.311) correlated positively with northerly aspect, although these latter two in a weaker manner. The second axis seemed to represent the differences in C, N, and particularly in C/N ratio (*r* = 0.998) among the sites as well as the varying dominance of hornbeam (*r* = -0.810) *versus* beech (*r* = 0.951) on the north-facing slopes. Richness of only two functional groups showed significant correlation with the NMDS ordinations: generalist saprotrophs correlating negatively with both the first (*r* = -0.835) and second axes (*r* = -0.551), while lichens predominantly correlated with the second axis (*r* = 0.934) and to a lesser extent with the first axis (*r* = -0.356). As for taxonomic groups with significant results, richness values of Dothideomycetes (*r* = -0.976), Eurotiomycetes (*r* = -0.848), and Sordariomycetes (*r* = -0.753) were negatively, while that of Mortierellomycota and Mucoromycota (*r* = 0.921) was positively correlated with northerly aspect (Fig. 3).

**Fig. 3.**
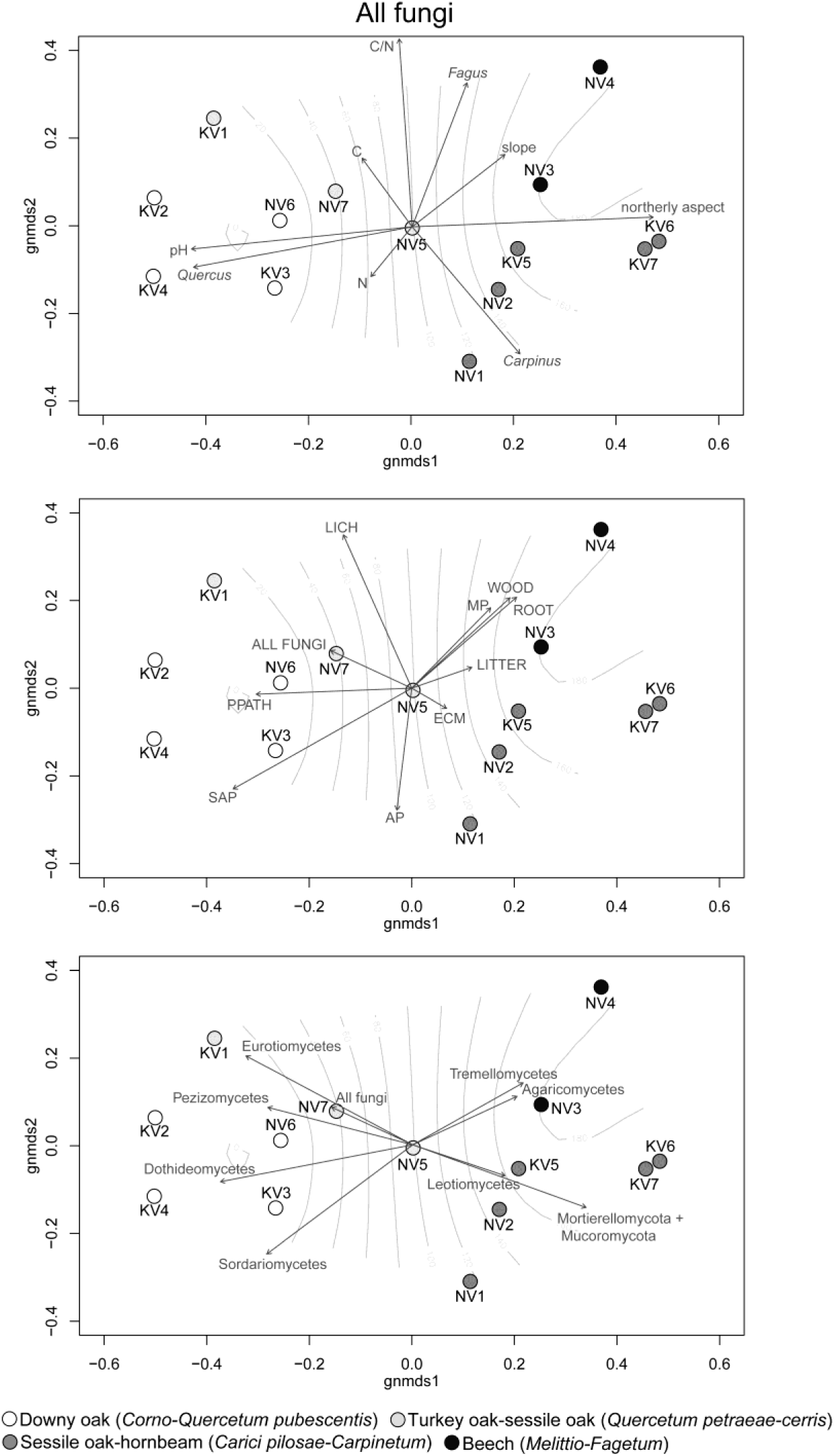
Non-metric multidimensional scaling (NMDS) ordination plots of the fungal communities in the sampled forest types based on Hellinger-transformed data, with northerly aspect displayed as isolines. Labels, localities and descriptions of the sampling sites are given in Table 1. Vectors of environmental variables and richness of functional and taxonomic groups of fungi correlated with ordination axes are displayed in three identical ordination plots. Abbreviations for functional guilds: AP = animal pathogen, ECM = ectomycorrhizal fungus, MP = mycoparasite, PPATH = plant pathogen, ROOT = root-associated (non-ECM) fungus, SAP = generalist saprotroph, WOOD = wood decomposer.

NMDS plots of the datasets corresponding to the functional groups of fungi showed similar correlations with aspect, forest types, and edaphic variables as detailed above. The NMDS ordinations of four largest functional groups in terms of OTU richness, i.e. ECM fungi, plant pathogens, generalist saprotrophs, and wood decomposers, are shown in Fig. 4, while those of the three less species-rich functional groups, i.e. animal pathogens, litter decomposers, mycoparasites, and root-associated fungi are shown in Fig. S1. Richness values of taxonomic families (or genera, when a family was represented by a single genus) of ECM fungi that correlated with northerly aspect were as follows: only ascomycete genera, i.e. *Elaphomyces*, *Helvella*, and *Tuber* showed negative correlations with northerly aspect, while only basidiomycetes, i.e. *Amanita*, Atheliaceae, *Cortinarius*, *Inocybe*, *Laccaria*, Russulaceae, and *Sebacina* correlated positively with northerly aspect. In addition, *Cenococcum*, *Clavulina*, and Thelephoraceae showed positive correlation with axis 2. Vectors representing the richness of taxonomic groups of plant pathogens seemed to be relatively evenly distributed in the ordination plot. Of these, Bionectriaceae and Mycosphaerellaceae showed preference for hornbeam forests, while *Leptodontitidum*, Taphrinaceae, and Venturiaceae seemed to favor beech forests, both on north-facing slopes. Conversely, richness in Coniochaetaceae, Didymellaceae, Massarinaceae, Phaeosphaeriaceae, *Pyrenochaeta*, and Spizellomycetaceae correlated strongly with southerly aspect. With regard to generalist saprotrophs, the majority of taxa showed clear preference for south-facing slopes, such as *Acremonium*, Aspergillaceae, Chaetomiaceae, Didymosphaeriaceae, Gomphaceae, Helotiaceae, Lasiosphaeriaceae, Microascaceae, Onygenaceae, Sporormiaceae, Stachyobotryaceae, *Tetracladium*, Trichocomaceae, Tricholomataceae, Trichomeriaceae, and Trichosporonaceae, while Mortierellaceae, Tremellaceae, Umbellopsidaceae, and to some extent Cunninghamellaceae tended to have higher richness in sites of northerly aspect. In wood decomposers, Corticiaceae, Hyaloscyphaceae, Lophiostomataceae, and *Pluteus* correlated negatively, while Chaetosphaeriaceae, Crepidotaceae, Helotiaceae, *Trechispora*, and Xylariaceae correlated positively with northerly aspect. MRPP confirmed the importance of aspect in shaping fungal community composition (effect size *A* = 0.084, probability *p* < 0.001). There were 137 and 174 significant (*p* < 0.05) indicator fungal OTUs characteristic of north-and south-facing slopes, of which, 46 and 69 were assigned to functional groups, respectively (Table 2).

**Table 2.**
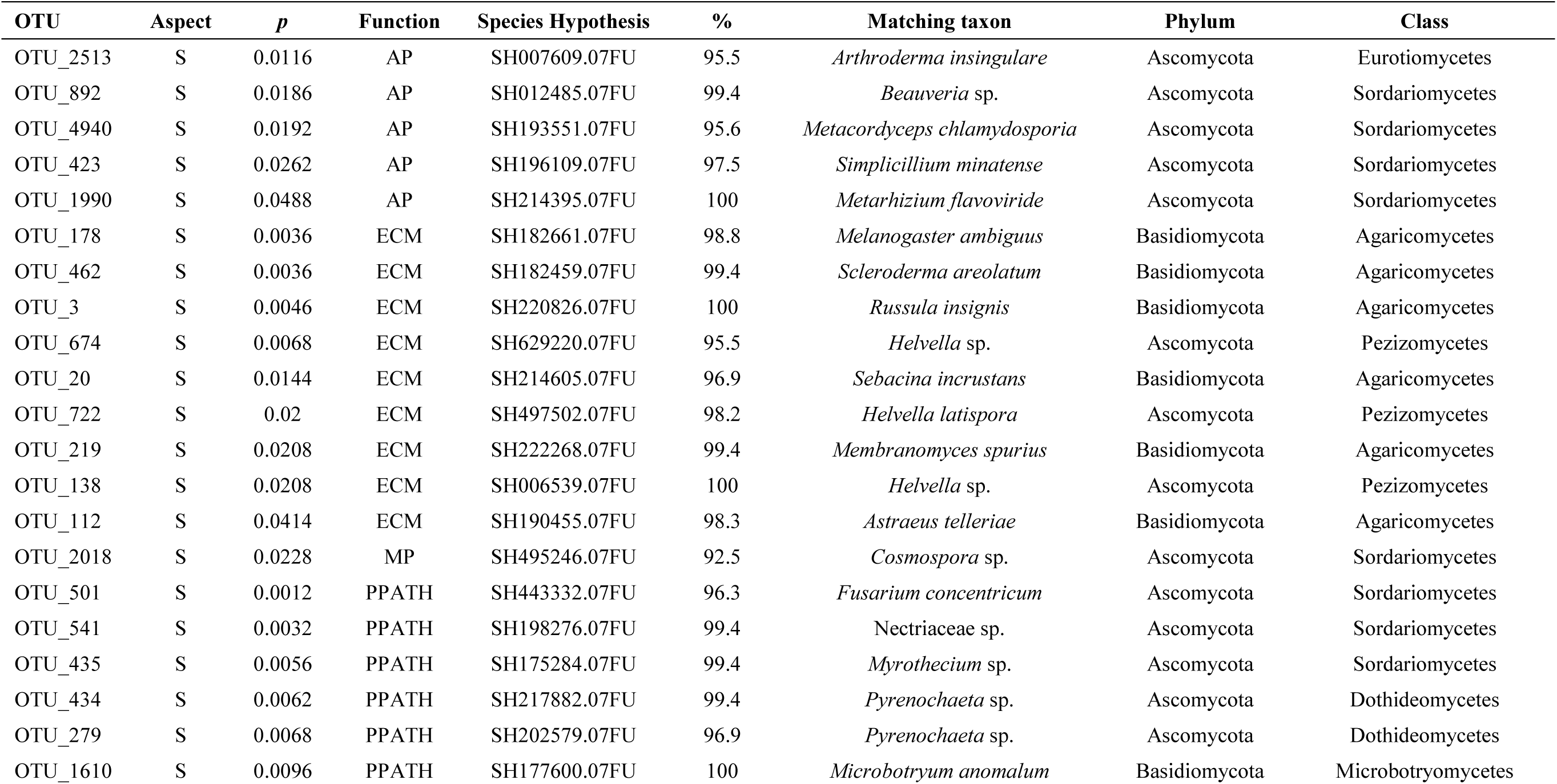
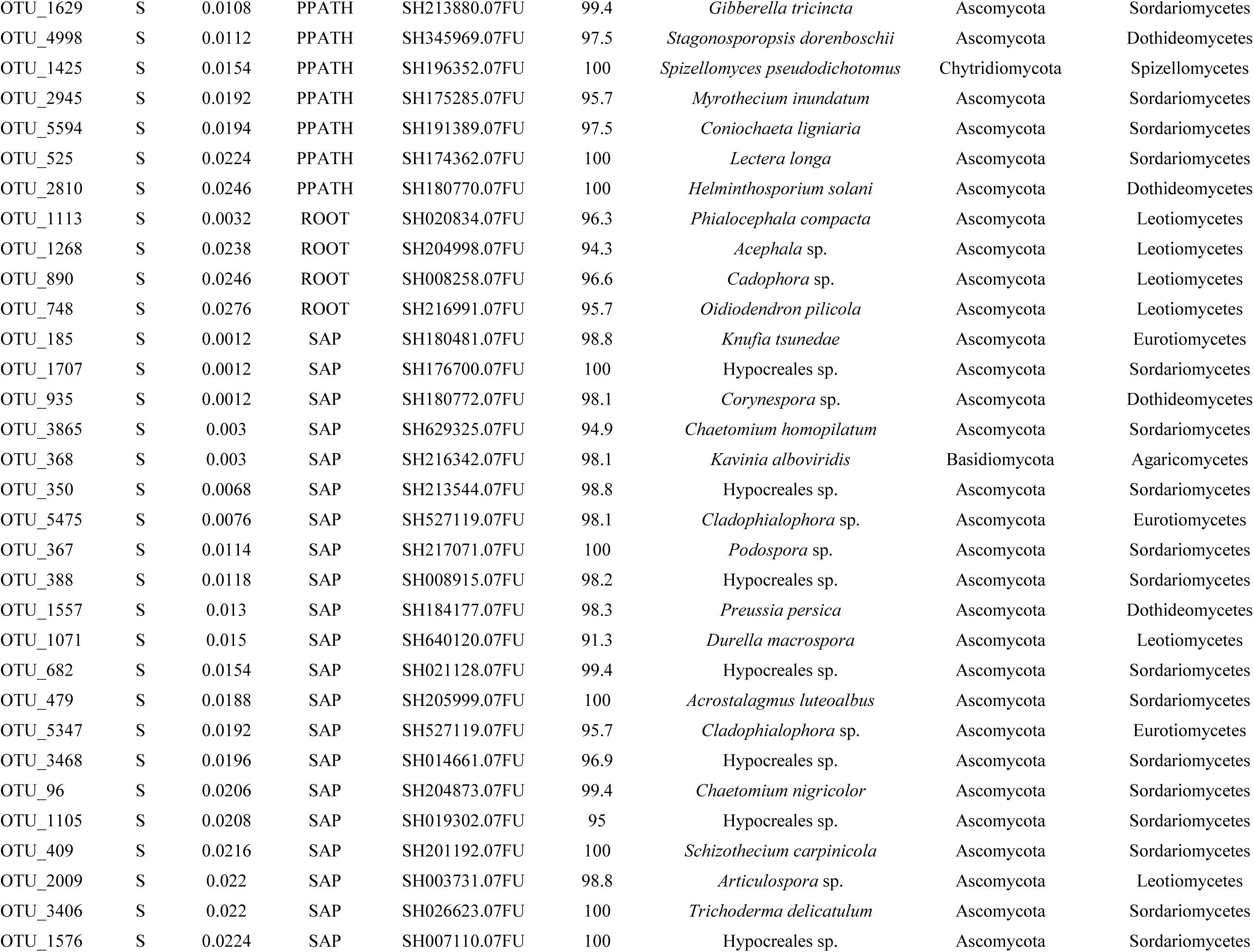
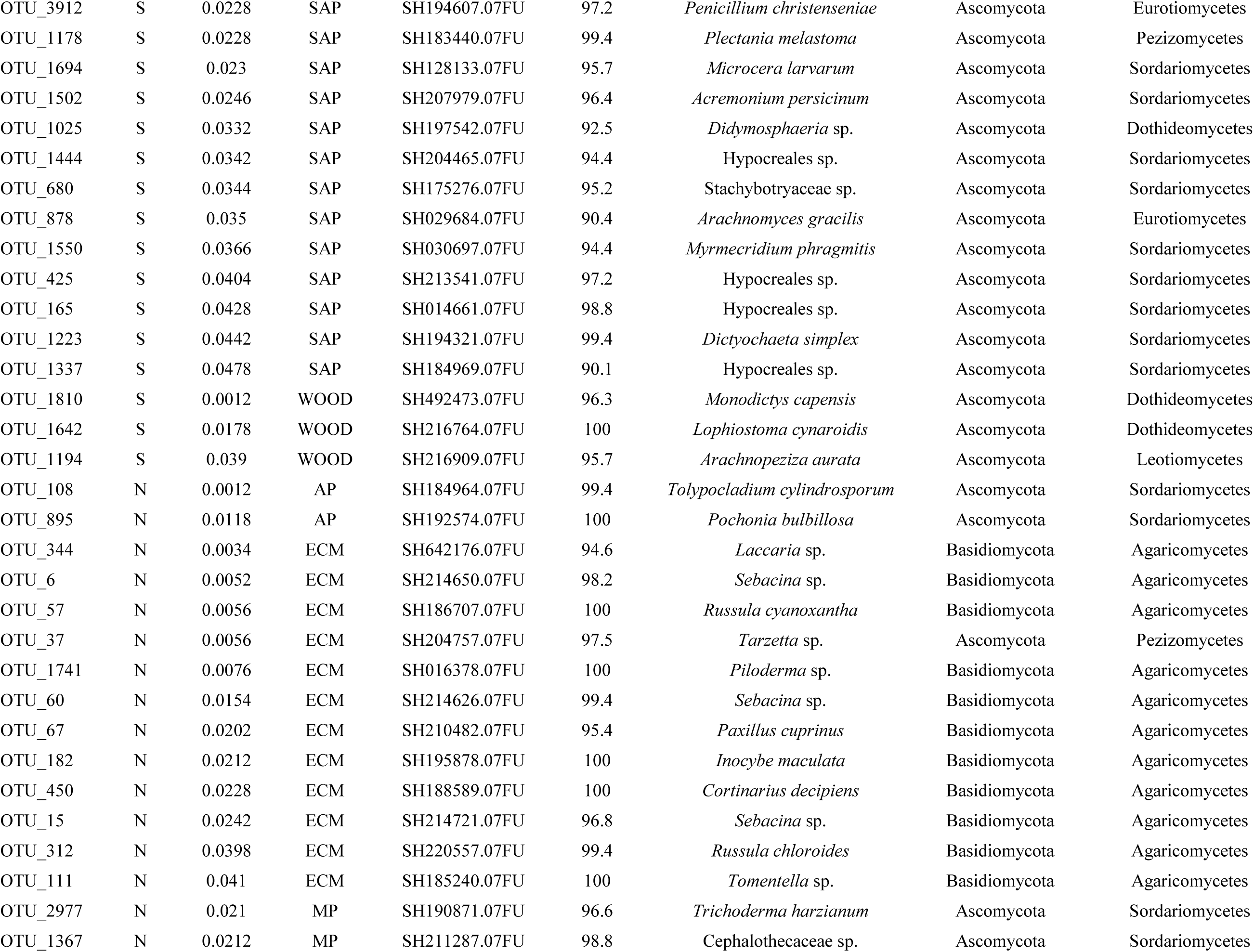
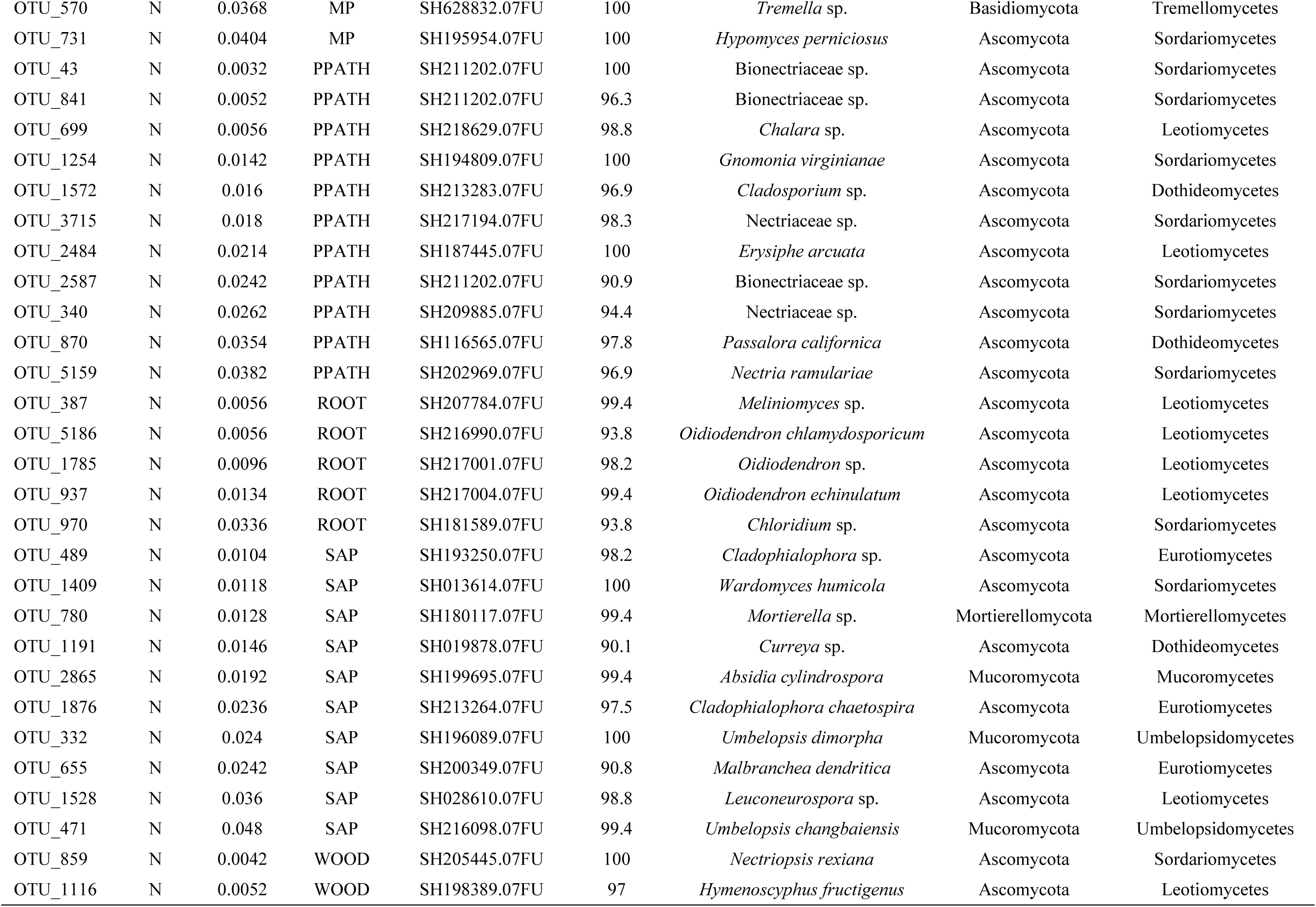
Fungal OTUs considered as significant indicators of northerly (N) or southerly (S) slope aspects with corresponding *p*-values, assigned functional guild, matching Species Hypothesis, ITS2 rDNA sequence similarity (%) and taxonomic classification of the most similar matching sequence in the UNITE+INSD dynamic Species Hypotheses database (version released on October 10, 2017). Only indicators with a known function are shown, displayed in the order of aspect, functional guild, and decreasing significance of indicator value. Abbreviations for functional guilds: AP = animal pathogen, ECM = ectomycorrhizal fungus, MP = mycoparasite, PPATH = plant pathogen, ROOT = root-associated (non-ECM) fungus, SAP = generalist saprotroph, WOOD = wood decomposer.

**Fig. 4.**
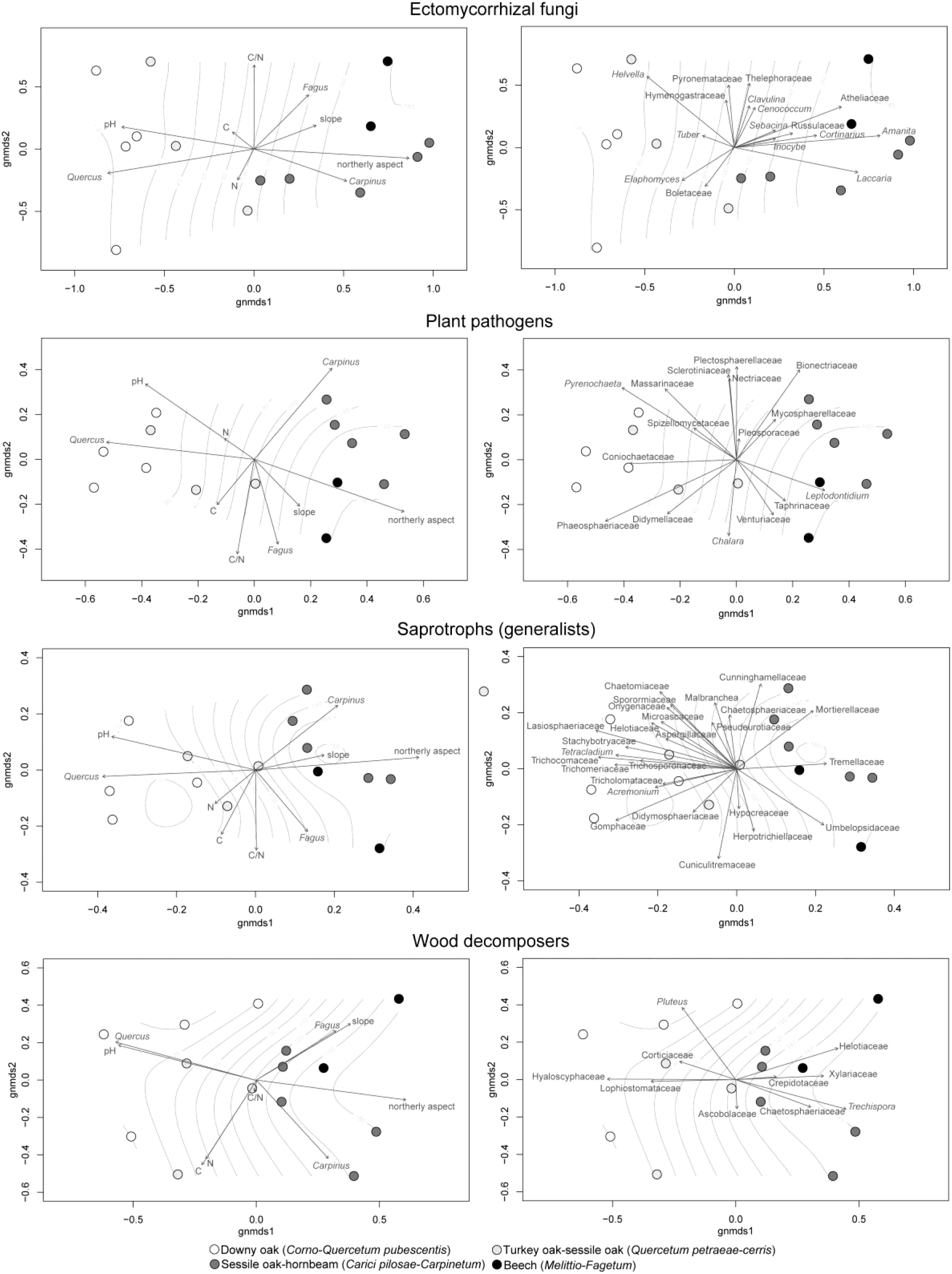
Non-metric multidimensional scaling (NMDS) ordination plots of the four more diverse functional groups of fungi in the sampled forest types based on Hellinger-transformed data, with northerly aspect displayed as isolines. Vectors of environmental variables and OTU richness of fungal families correlated with ordination axes are displayed. Where a certain family was represented by only one genus, the genus name is shown.

### Assessing the effects of environmental factors on fungal community composition

Mantel tests showed that neither spatial proximity nor slope had any significant correlation with fungal community composition or with aspect, relative abundance of tree genera, and edaphic factors. On the other hand, aspect was strongly correlated with the relative abundance of tree genera (*r =* 0.526; *p <* 0.001) and weakly with edaphic factors (*r =* 0.151; *p =* 0.046), while there was moderate correlation between tree genera and edaphic factors (*r =* 0.266; *p <* 0.035) (Table 3). Fungal community composition was strongly correlated with aspect (*r* = 0.743, *p* < 0.001), relative abundance of tree genera (*r* = 0.558, *p* < 0.001), and edaphic factors (*r* = 0.535, *p* < 0.001). Nonetheless, partial mantel tests indicated that aspect in itself had a strongly significant effect on community structure (*r* = 0.637, *p* < 0.001) when tree genera was accounted for (control matrix), while the effect of tree genera was substantially weaker, though still significant (*r* = 0.295, *p* = 0.018) when aspect was controlled. Conversely, the correlations of aspect (*r* = 0.793, *p* < 0.001) and edaphic factors (*r* = 0.639, *p* < 0.001) with fungal community composition were both strong when edaphic factors or aspect were controlled for, respectively (Table 3).

**Table 3.** Correlation of environmental and spatial variables with each other and with fungal community composition of the sampling sites. Correlations among standardized environmental variables and with Hellinger-transformed fungal community matrix were tested individually using Mantel tests showing correlation coefficients (*r*) and significance values (*p*). Non-significant results (n.s.) are not shown. In addition, to disentangle the ‘pure’ effects of environmental variables on fungal community composition, partial Mantel tests were used in all combinations of tested and controlled environmental variables.

| Correlation among environmental variables |  |  |  |  |
| --- | --- | --- | --- | --- |
|  | aspect | slope | tree genera | edaphic factors |
| aspect | - |  |  |  |
| slope | n.s. | - |  |  |
| tree genera | $r=0.5256; p<0.0001$ | n.s. | - | |
| edaphic factors | $r=0.1508; p=0.0461$ | n.s. | $r=0.2661; p=0.0354$ | - |
| spatial | n.s. | n.s. | n.s. | n.s. |

| Correlation with fungal community composition |  |  |  |  |  |  |
| --- | --- | --- | --- | --- | --- | --- |
| Tested variables | Mantel test | Control variables for partial mantel test |  |  |  |  |
|  |  | aspect | slope | tree genera | edaphic factors | spatial |
| aspect | $r=0.7429; p<0.0001$ | - | $r=0.7395; p<0.0001$ | $r=0.6368; p<0.0001$ | $r=0.7928; p<0.0001$ | $r=0.7449; p<0.0001$ |
| slope | n.s. | n.s. | - | n.s. | n.s. | n.s. |
| tree genera | $r=0.5584; p<0.0001$ | $r=0.2949; p=0.0178$ | $r=0.5486; p=0.0002$ | - | $r=0.5108; p=0.0002$ | $r=0.5579; p=0.0001$ |
| edaphic factors | $r=0.5347; p<0.0001$ | $r=0.6388; p<0.0001$ | $r=0.5247; p=0.0002$ | $r=0.4828; p=0.0006$ | - | $r=0.5397; p<0.0001$ |
| spatial | n.s. | n.s. | n.s. | n.s. | n.s. | - |

Variation partitioning analyses indicated that the tested environmental variables explained 24.3% of the total variation observed. Topography (aspect and slope) explained 15.1% of the variation, but only 0.8% after accounting for the relative abundance of dominant tree genera and edaphic factors. Tree genera accounted for 14.7% of the variation and 3.1% when the effect of other variables was removed. Measured soil variables explained 17.9% of the variation, of which 5.9% was attributed to their ’pure’ effect. The shared variation explained by topography and tree genera and by topography and edaphic factors was 2.5% and 2.9%, respectively, while the shared effects of all three groups of variables accounted for 8.9% of the total variation (Fig. S2).

## 4. DISCUSSION

### Drivers of landscape-level compositional patterns of fungal communities

The deep sequence data presented here clearly show that fungal community composition at the selected Pannonian forest sites is strongly structured according to slope aspect. Although some fungal species occurred in all samples, the majority of fungal OTUs preferred either south-or north-facing slopes, as suggested by the observed community turnover as well as the numerous indicator species. Similarly, even though total fungal richness was not statistically different between north-and south-facing slopes, many functional and taxonomic groups of fungi were more diverse on slopes of northerly or southerly aspect.

Naturally, the effect of slope aspect on fungal community composition and richness is mediated through abiotic and biotic factors that are driven either directly or indirectly by the differences in net solar radiation received in north-and south-facing slopes. For example, aspect strongly influences local air and surface temperature, soil moisture, relative humidity, and soil chemical processes (Gilliam *et al.* 2014). Consequently, the habitats found on north-and south-facing slopes have distinct meso-and microclimatic as well as edaphic conditions. These differences are particularly marked regarding available moisture, soil organic matter, and pH, as had been observed in the study region previously (Dobos 2010), which, in turn, influence the composition of biotic communities and their interactions. Although the relative abundance of the two dominant tree genera (*Carpinus* and *Quercus*) was strongly influenced by aspect, they occurred in most of the sites. Therefore, and because the vast majority of plant-associated fungi are not strictly specific to a tree genus or family, I argue that to a great extent the observed fungal community patterns are caused by a complex array of aspect-driven environmental variables and not by the type of vegetation alone.

The availability of soil moisture is one of the most important environmental conditions that determines richness as well as community composition in fungi (Crowther *et al*. 2014; Tedersoo et al. 2014). Because at landscape scale, as in the study area, annual precipitation is considered uniform across the sampling sites due to their close proximity, topography likely is the most important factor that influences moisture availability through contrasting levels of evapotranspiration between north-and south-facing slopes, as has been observed in several biomes (Fekedulegn *et al*. 2003; Méndez-Toribio *et al*. 2016). Drought-tolerant fungi are generally regarded as generalists, instead of dry specialists, with respect to their ability to grow along a broad moisture gradient (Lennon *et al*. 2012), while drought-intolerant fungi are considered specialists to a narrow range of mesic conditions, where they likely have competitive advantage over the generalists (Crowther *et al*. 2014). The resulting competitive dynamics may force drought-tolerant fungi to the more xeric south-facing slopes, while fungi with a stronger competitive ability under mesic conditions are expected to dominate the mesophilous forests on the north-facing slopes in the study area.

Soil pH is also known to play an important role in shaping fungal communities (Porter *et al.* 1987; Coughlan *et al.* 2000; Lauber *et al.* 2008; Rousk *et al.* 2010; Geml *et al*. 2014a; Tedersoo *et al*. 2014; Glassman *et al.* 2017) and is often influenced by slope aspect (Gilliam *et al*. 2014; Chu *et al.* 2016). Because many fungal species have a relatively wide pH optimum (e.g. Wheeler *et al.* 1991; Nevarez *et al.* 2009), it is likely that the observed correlation of pH with community composition is mainly indirect, e.g., via altering nutrient availability and competitive interactions between soil fungi and bacteria (Rousk *et al.* 2008), and other soil biota. In the study sites, there was a strong negative correlation of soil pH with northerly aspect and the interaction of topography (mainly aspect) and edaphic factors (primarily pH) provided a large fraction of the explained variation in fungal community composition. Therefore, it is difficult to disentangle the ‘pure’ effect of pH from that of aspect. Nevertheless, for some fungal groups that are known to be influenced by soil pH, the data presented here are in agreement with previous results. For example, root endophytic fungi had been shown to prefer low soil pH (Postma *et al.* 2006) and the strong preference of non-ECM root-associated fungi for the northerly sites with lower pH in this study confirms the above trend Similarly, ECM fungi are generally considered acidophilus (Read 1991), and there was weak, although non-significant, positive correlation between ECM fungal richness and northerly aspect (i.e. sites with lower pH). The only other edaphic factors with strong correlation with fungal community structure was C/N ratio, which was not related to aspect. Instead, changes in C/N ratio appeared to be related to different forest types on the north-facing slopes, i.e. oak-hornbeam and submontane beech forests. Because C/N is considered a direct measure of resource quality (Nielsen *et al*. 2010), it is possible that the higher C/N values in the beech forests are driven by differences in litter quality between beech and hornbeam and oak. Measurements from more beech and oak-hornbeam stands are needed to test this htpothesis.

### The contrasting effects of aspect on functional groups of fungi

The data clearly show a strong emerging pattern driven by slope aspect in all fungal groups. Several functional groups showed strong differences in richness between north-and south-facing slopes. Mycoparasites, non-ECM root-associated fungi, and wood decomposers had higher richness in the sites with predominantly northerly aspect. The cooler microclimate in particular may be more advantageous for root-associated fungi, because this group has been shown to be more diverse at higher elevations in altitudinal gradient studies (Geml *et al*. 2014b). On the other hand, although richness of wood decay fungi and mycoparasites have been shown to decline with decreasing temperature in elevation gradients (Geml *et al*. 2014b), considering the given moisture limitation in this study area, their higher richness in north-facing slopes may be more strongly driven by the greater availability of moisture. Lichens, plant pathogens and generalist saprotrophs were clearly more species rich in the south-facing slopes. With regard to lichens, this may be a consequence of the more open canopy of the thermophilous oak forests that allow more light to penetrate and apparently fosters the growth of the generally shade-intolerant and drought-tolerant lichens. On the other hand, plant pathogens and generalist saprotrophs may benefit more from the higher temperatures characteristic of the south-facing slopes, as richness values of these groups generally correlate positively with temperature in altitudinal as well as in global studies (Geml *et al*. 2014b, Tedersoo *et al*. 2014). Also, many generalist saprotrophs (e.g., in Eurotiomycetes) are known for their drought-tolerance and, based on the above-mentioned theory on competitive dynamics, they may be outcompeted in the mesic sites by mesophilic specialists, resulting in their higher diversity in the warmer and drier south-facing slopes.

Even in functional groups with no significant differences in richness, I observed strong compositional differences between slopes of northerly and southerly aspects. For example, in ECM fungi, ascomycete and basidiomycete genera were clearly more diverse in the south-and north-facing slopes, respectively. This is in agreement with other studies showing that ECM communities often are dominated by ascomycetes in arid and semiarid environments, while basidiomycetes tend to dominate more moist habitats (Cavender-Bares *et al*. 2009; Gehring *et al*. 2014). This may be linked to their foraging and fruiting strategies and the related differences in C requirement from the host trees. For example, trees in more xeric conditions are expected to preferentially associate with ECM fungi that have contact and short-distance, as opposed to medium-and long-distance, extrametrical mycelial exploration types and smaller fruiting bodies that are less costly in terms of C requirement.

### The unexplained component of community turnover

An ever-present feature of fungal community studies is the large amount of unexplained variation in richness and community composition even at small spatial scales (Peay *et al*. 2016). In this study, the most significant compositional differences were observed between slopes of northerly and southerly aspects, but there was also substantial variation among sites within sites on the same slope. The environmental variables measured in this study explained about one quarter of the variation in fungal community composition in all sites, which confirms the above-mentioned substantial unexplained component of community assembly. Because many OTUs were rare, i.e. found only in one or two sites, most such differences may be due to random processes in community assembly as well as due to random sampling, as truly exhaustive soil sampling is practically impossible to achieve in the field. The former is partly explained by the priority effect, i.e. within a given species pool of a particular habitat, stochastic dispersal determines the order in which newly available resources are colonized by different species, which, in turn, drives to a large extent the composition of the community (Peay *et al.* 2016). In addition, other factors not examined here, such as density-dependent processes (e.g., intra-and interspecific competitions and pathogen-host interactions) as well potentially important environmental variables not yet measured, may also contribute to the observed beta diversity. For example, I did not measure soil and air temperature, relative humidity, and soil moisture at the sites, partly because it is already well-known that these factors correlate strongly with aspect on mesoclimate scale, as mentioned above, and partly because obtaining a realistic characterization of these variables would have required measurements taken throughout the growing season at the sampling sites, which was beyond the scope and logistic possibilities of this case study. However, by including site-specific microclimate data as well as a more extensive list of edaphic variables, future studies may obtain more insights into the variation of environmental factors at small spatial scales as well as their influence on fungal community composition and turnover.

### Contribution to the knowledge on Pannonian forest fungi

This study shows a remarkably high fungal diversity in a small area (< 2 km^2^) of secondary forests in the eastern edge of the Mátra mountains. After the stringent quality filtering steps, the non-rarefied and rarefied datasets contained 4695 and 4312 fungal OTUs, respectively. Based on the rarefied data, well over 1000 OTUs occurred in any given sample (Fig. 2), each representing an area of ca. 250 m^2^. In total, I detected representatives of 707 fungal genera, of which 467 belonged to Ascomycota, 225 to Basidiomycota, 6 to Chytridiomycota, and 7 represented early-diverging lineages formerly classified in Zygomycota. Nonetheless, the true generic diversity likely is even higher, because the vast majority of fungi in the sampled sites, as well as globally, are microfungi and many of them could only be assigned to families, orders or classes due to the lack of sufficiently identified reference data. Consequently, these represent species with unknown identity, several of which may still be undescribed. With respect to the identified microfungi, the results presented here may be the first data on their diversity and possible habitat preference in the Pannonian biogeographic region, serving as potential reference data for future studies as well. By providing the full list of taxa corresponding to the 2542 unique SHs that matched the OTUs in these samples with high (> 95%) sequence similarity (Table S1), I intend to facilitate other mycological and fungal ecological studies in the region.

I was able to assign a relatively high proportion (51%) of fungal OTUs to functional groups, particularly in taxa with macroscopic fruiting bodies for which extensive reference data are available from Europe, e.g., agarics, boletes, coral fungi, polypores, and several true and false truffles. For example, the number of unique matching SHs in some of the most diverse basidiomycete genera were 69 in *Inocybe*, 48 in *Russula*, 18 in *Lepiota*, while somewhat surprisingly *Cortinarius* and *Lactarius* were only represented by 23 and 13 unique SHs, respectively. On the other hand, the data also showed a high diversity ECM basidiomycetes with inconspicuous fruiting bodies, such *Sebacina* (66 SHs) and *Tomentella* (83 SHs) that are notoriously underrepresented in sporocarp surveys (Gardes & Bruns 1996; Kõljalg *et al*. 2000; Geml *et al*. 2012). Furthermore, I detected numerous hypogeous ECM fungi, such as *Elaphomyces muricatus* and *E. papillatus*, *Genea verrucosa*, *Hydnotrya* sp., *Hymenogaster australis*, *H. griseus*, *Hysterangium stoloniferum*, *Gautieria graveolens*, *Melanogaster ambiguus*, *M. broomeanus*, and *M. variegatus*, *Wakefieldia macrospora*, as well as several species of the charismatic *Tuber* genus (e.g., *T. aestivum*, *T. borchii*, *T. brumale*, *T. maculatum*, *T. puberulum*, *T. rapaeodorum*, *T. rufum* and possibly *T. fulgens*). Most of these are known to occur in Hungary, except for *Gautieria graveolens*, as this genus is only represented by an unidentified species in the list of hypogeous fungi for the Carpathian-Pannonian region compiled by Bratek *et al.* (2013). Representatives of these hypogeous fungi were found in all forest types, indicating that they probably are relatively common in Pannonian forests.

In addition, I want to emphasize the suitability of DNA metabarcoding to complement sporocarp-based assessments for biological monitoring and conservation of fungi. Specifically, studies such as this can provide valuable knowledge not only on the total diversity of fungi at any given site, but also on the distribution and habitat preferences of numerous species, including rare and/or protected species. For example, I found at least one of the thirty-five species of fungi protected by law in Hungary (Siller *et al*. 2005, 2006): *Strobilomyces strobilaceus*, which was detected on north-facing slopes of both the Kis-and Nagy-Várhegy, which confirms the habitat preference of this species for mesophilic beech and oak-hornbeam forests in Hungary (Siller *et al.* 2005). Furthermore, even though the retained global singletons that were > 98% similar to a reference fungal SH had only minor contribution to richness values and had practically no influence on the community structure patterns, 238 were identified to a genus or species. Of these, the following taxa were only represented by singletons in the dataset: *Aleurodiscus aurantiacus*, *Annulohypoxylon multiforme*, *Aspergillus cibarius*, *Baeospora myosur*a, *Barbatosphaeria dryina*, *Bullera alba*, *Calonectria quinqueseptata*, *Caloplaca obscurella*, *Chaetosphaeria decastyla*, *Chroogomphus mediterraneus*, *Clavariadelphus* sp., *Coprinopsis picacea*, *Coprinus comatus*, *Cortinarius casimiri*, *Cortinarius cotoneus*, *Cortinarius infractus*, *Cortinarius pardinus*, *Crepidotus mollis*, *Cryptodiscus pallidus*, *Cylindrocladiella elegans*, *Dactylellina ellipsospora*, *Darksidea epsilon*, *Durella connivens*, *Elaphomyces papillatus*, *Elmerina caryae*, *Exidia truncata*, *Filobasidium magnum*, *Geastrum fornicatum*, *Geopora* sp., *Gloeocystidiellum kenyense*, *Gromoniopsis idaeicola*, *Haptocillium campanulatum*, *Hyalorbilia erythrostigma*, *Hypogymnia physodes*, *Jattaea tumidula*, *Kretzschmaria deusta*, *Lactarius necator*, *Leiosepium tulasneanum*, *Leptosphaeria rubefaciens*, *Leptospora rubella*, *Leucoagaricus barssii*, *Lophiotrema eburnoides*, *Melanelixia subaurifera*, *Melanospora damnosa*, *Microstroma phylloplanum*, *Mycena renati*, *Mycosphaerella ulmi*, *Neocatelunostroma abietis*, *Neocladophialophora quercina*, *Ossicaulis lignatilis*, *Otidea onotica*, *Paxillus obscurisporus*, *Peltaster fruticola*, *Peniophorella praetermissa*, *Pestalotiposis chinensis*, *Phlebia tremellosa*, *Pholiotina teneroides*, *Phyllozyma linderae*, *Pilidium concavum*, *Pleurophoma ossicola*, *Pluteus velutinus*, *Psilocybe inquilina*, *Resupinatus applicatus*, *Ramphoria pyriformis*, *Rhytisma acerinum*, *Russula albonigra*, *Russula inamoena*, *Russula sororia*, *Sarcosphaera coronaria*, *Scheffersomyces stipitis*, *Sistotrema brinkmannii*, *Slopeiomyces cylindrosporus*, *Sphaerollopsis macroconidialis*, *Steccherinum ochraceum*, *Stylonectria norvegica*, *Thielavia subthermophila*, *Tomentella badia*, *Trechispora laevis*, *Tremella yokohamensis*, *Tricholoma batschi*, *Urocystis agropyri*, *Ustilago nunavutica*, *Valsivia insitiva*, *Vanrija humicola*, *Verrucaria dolosa*, *Xenasmatella borealis*, *Xylodon rimossimus*, *Xylomelasma* sp., *Zalerion arboricola*, and *Zasmidium dalbergiae*. Most of the macrofungi among these, such as *Baeospora myosur*a, *Coprinopsis picacea*, *Coprinus comatus*, *Cortinarius infractus*, *Crepidotus mollis*, *Geastrum fornicatum*, *Mycena renati*, *Russula albonigra*, *Russula heterophylla*, *Russula sororia*, and *Tricholoma batschi*, have been found in sporocarp surveys in the Északi-középhegység (Bohus & Babos 1960; Tóth 1999; Siller *et al.* 2002; Egri 2007; Rudolf *et al.* 2008; Siller & Dima 2014). With respect to the rest, more sampling is needed to confirm their presence in the region.

Finally, soil communities are extremely diverse and there is increasing evidence pointing to soil biodiversity as having key roles in determining the structure and ecological responses of terrestrial ecosystems (Bardgett & van der Putten 2014). Soil fungi in particular are known to drive plant diversity and productivity and are crucial for ecosystem functioning and resilience towards disturbance (van der Heijden *et al.* 2008). Because most fungi have high habitat specificity and tend to respond quickly to changes in environmental conditions (Nielsen *et al.* 2010; Geml *et al*. 2015, 2016; Morgado *et al.* 2015, 2016; Mundra *et al.* 2016; Semenova *et al*. 2015, 2016), fungi have a promising potential as indicators of habitat quality in biological monitoring programs. More specifically, assessments of richness and community composition of fungal communities in a variety of habitats can inform decision-makers with respect to land use strategies that foster the sustainable preservation of diverse and resilient ecosystems with a wide range of ecosystem functions.

### Conclusions

A prominent finding of this study is that fungal diversity and community structure are strongly influenced by slope aspect, despite the short distance separating north-and south-facing slopes. Even though this finding may appear trivial due to the well-known effects of aspect on vegetation, fungal communities are surprisingly little studied in this regard and the data presented here offer unprecedented insights into the landscape-level distribution of various taxonomic and functional groups with respect to topography. Furthermore, while aspect-driven differences in vegetation tend to be related to relative importance as opposed the presence/absence of plant species on slopes with different aspect (Gilliam et al. 2014), many fungal species were detected exclusively on either north-or south-facing slopes. This reflects the often-observed high habitat specificity exhibited by many fungi, which offers possibilities for biological monitoring and habitat characterization and I strongly advocate for incorporating fungi in biodiversity assessments and conservation efforts.

## ACKNOWLEDGEMENTS

I am grateful to Péter Molnár for his assistance during the fieldwork, to Marcel Eurlings (Naturalis) for carrying out the Ion Torrent sequencing, to Luis Morgado (University of Oslo and Naturalis) for sharing the R script for the NMDS analysis, to Eszter Draskovits and Nikolett Tarjányi (Institute for Soil Sciences and Agricultural Chemistry, Hungarian Academy of Sciences, Budapest) for their assistance in obtaining the soil chemical data, to József Sulyok and András Schmotzer (Bükk National Park) for sending maps and aerial photographs of the sampling region, and to Attila Baranyi for preparing the map for Fig. 1. I also thank Jeremy Miller (Naturalis) for his constructive comments on the manuscript. The molecular work and the soil chemical analyses were supported by the Naturalis Research Initiative fund provided to József Geml.

---

**Fig. S1.**
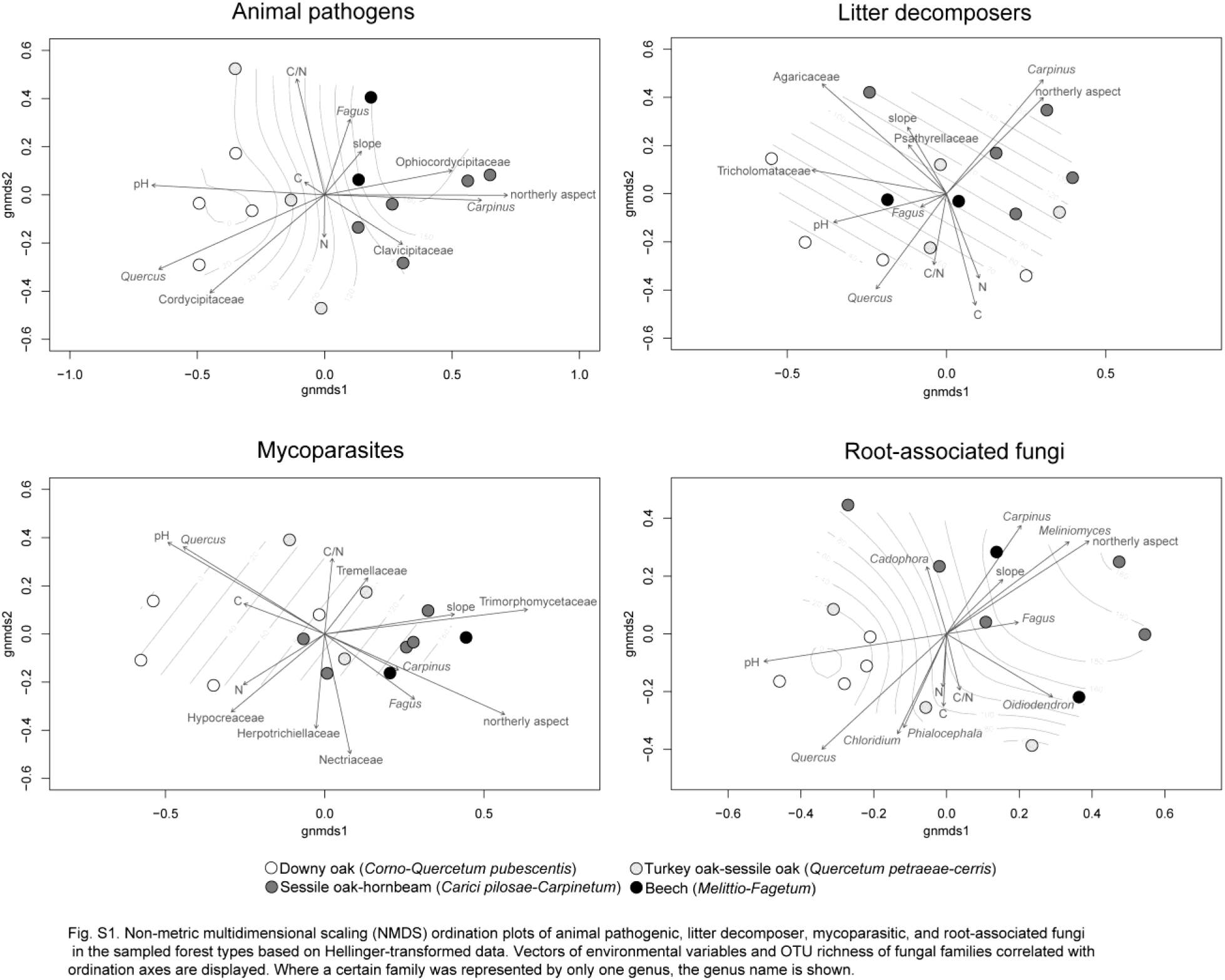
Non-metric multidimensional scaling (NMDS) ordination plots of animal pathogenic, litter decomposer, mycoparasitic, and root-associated fungi in the sampled forest types based on Hellinger-transformed data. Vectors of environmental variables and OTU richness of fungal families correlated with ordination axes are displayed. Where a certain family was represented by only one genus, the genus name is shown.

**Fig. S2.**
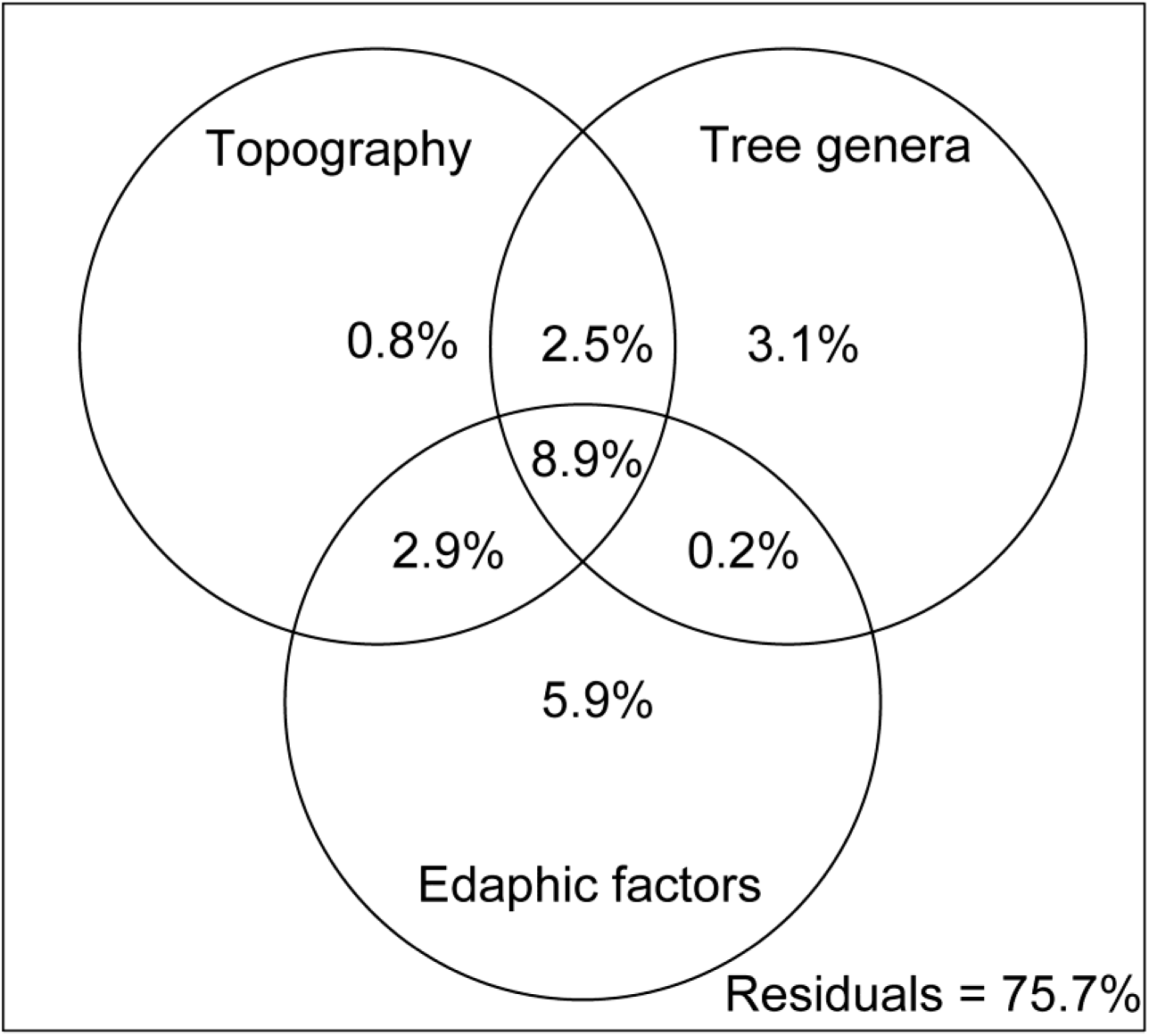
The contribution of environmental variables to explaining the variation in fungal community composition among the sampling sites as estimated by variation partitioning analyses. Tested environmental variables included topography (aspect, slope), relative abundance of tree genera (*Carpinus*, *Fagus*, *Quercus*), and edaphic factors (soil pH, C, N, and C/N)

**Table S1.** The full list of taxa corresponding to the 2542 unique SHs that matched the OTUs in the collected samples with high (> 95%) sequence similarity, with % ITS2 rDNA sequence similarity to the most similar OTU, Species Hypothesis code, and taxonomic classification. Taxa are ordered according to phylum, class, order, family, genus, and the identified taxon name (provided upon manuscript acceptance).

